# The neuroprotective effect of meloxicam in a transient ischemia model involves an increase in axonal sprouting but a decrease in new neuron formation after 7 days of reperfusion

**DOI:** 10.1101/2021.04.05.438505

**Authors:** IF Ugidos, P González-Rodríguez, M Santos-Galdiano, E Font-Belmonte, B Anuncibay-Soto, D Pérez-Rodríguez, A Fernández-López

## Abstract

The inflammatory response plays an important role in neuroprotection and regeneration after ischemic insult. The use of non-steroidal anti-inflammatory drugs has been a matter of debate as to whether they have beneficial or detrimental effects. In this context, the effects of the anti-inflammatory agent meloxicam have been scarcely documented after stroke, but its ability to inhibit both cyclooxygenase isoforms (1 and 2) could be a promising strategy to modulate post-ischemic inflammation. This study analyzed the effect of the anti-inflammatory agent meloxicam in a transient focal ischemia model in rats, measuring its neuroprotective effect after 48 hours and 7 days of reperfusion and the effects of the treatment on the glial scar and regenerative events such as the generation of new progenitors in the subventricular zone and axonal sprouting at the edge of the damaged area. We show that meloxicam’s neuroprotective effects remained after 7 days of reperfusion even if its administration was restricted to the two first days after ischemia. Moreover, meloxicam treatment modulated glial scar reactivity, which matched with an increase in axonal sprouting. However, this treatment decreased the formation of neuronal progenitor cells. This study discusses the dual role of anti-inflammatory treatments after stroke and encourages the careful analysis of both the neuroprotective and the regenerative effects in preclinical studies.

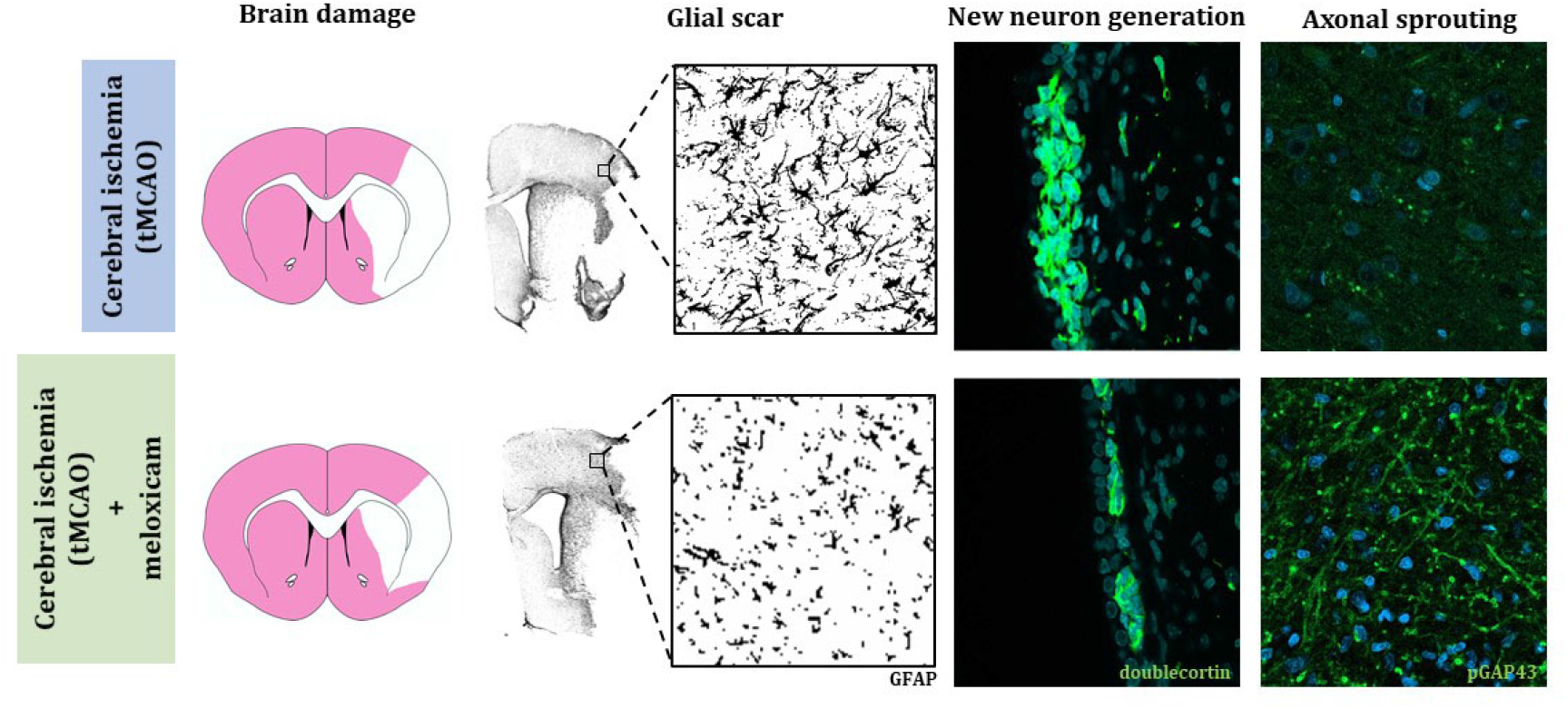

## 1. Introduction

Inflammation plays a major role in many central nervous system diseases and a crucial role in neuroprotection and regeneration after stroke [1–3]. Post-ischemic inflammation is triggered by free radicals and damage-associated molecular patterns produced by dying neurons [4,5] and involves the production of cytokines, the activation of microglia and astroglia cells, the infiltration of peripheral immune cells into the brain and, in later events, the formation of a glial scar surrounding the damaged area. These events activate positive pro-inflammatory feedback that exacerbates tissue damage [6,7]. Moreover, inflammation also plays a role in neurogenesis and neuronal repair after stroke [8,9]. The interaction between inflammation and the regenerative response has been widely studied, and whether it is beneficial or detrimental is still a matter of debate [10,11]. The extensive role of the inflammatory process in early and late events after stroke has led to the search for agents that modulate this pro-inflammatory loop as a putative palliative and regenerative strategy [12].

A key step in the inflammatory response is the activation of cyclooxygenases (COX). The isoform COX-2 has been historically targeted because it is highly inducible after stroke, compared to the isoform COX-1, which is considered to be constitutively expressed [5,7,13]. Moreover, excessive COX-1 inhibition has been related to peripheral side effects [14,15], driving pharmaceutical companies to develop selective COX-2 inhibitors. In this regard, the use of non-steroidal anti-inflammatory drugs (NSAIDs) that block either COX-1, COX-2, or both of them to a various extent has been widely discussed. NSAIDs are not expensive and their kinetics have been very well characterized. However, their effects after stroke are controversial. The general belief that the highly COX-2 selective blockers would improve neuroprotection resulted in controversy when several studies showed a detrimental effect in different models of ischemia [16–18], leading to the reconsideration of both the role of COX-1 inhibition in post-ischemic treatments [19– 21] as well as the proper balance of the inhibition of both isoforms.

Meloxicam is a preferential COX-2 inhibitor that exhibits COX-2 specificity between 3 and 77 times that of COX-1 [22,23] and is widely used in the veterinary clinic [24–26], as well as in humans [27,28]. Meloxicam is widely used as a post-operative treatment to reduce inflammation and pain with very efficient results [29,30]. Despite the wide use of this anti-inflammatory agent in the clinic, and some studies showing the positive effects of meloxicam [31,32], the effects of meloxicam as a palliative agent in the focal cerebral ischemia model is scarcely documented and it is mainly focused on their use as analgesic rather than in its neuroprotective effect [33]. This study shows, for the first time, short- and long-term effects of meloxicam in a model of focal cerebral ischemia by transient middle cerebral artery occlusion (tMCAO) and its interactions with regenerative events.

## 2. Material and methods

### 2.1. Animals

Sixty-four eight-week-old male Sprague-Dawley rats (320-360g) (Janvier Labs, Le Genest-Saint-Isle, France) were used to perform this study. Animals were housed at standard temperature (22 ± 1°C) in a 12 h light/dark cycle with food (Panlab, Barcelona, Spain) and water *ad libitum*. Animals were caged in pairs and randomly designated to an experimental group by a different person than those who performed the surgery.

The study was designed using six animals in each experimental group (Sham, Vehicle, Mel0.5, Mel1, Mel5, Mel10) to perform dose-response assays, behavioral tests, and protein and serum analysis at 48 hours of reperfusion. For immunofluorescence assays at 48 hours, four and five animals were used in each experimental group (Vehicle and Mel1, respectively) were used. At 7 days of reperfusion, five animals in each experimental group (Vehicle, Mel1-a, Mel1-c) were used to perform immunofluorescence and behavioral analysis. Five animals died during surgery. Three animals were discarded due to a lack of reperfusion after surgery. Four animals were also discarded because of insufficient blood flow restriction, and two animals were culled due to poor outcomes after stroke.

All procedures were carried out in compliance with the Animal Research: Reporting of In Vivo Experiments (ARRIVE) guidelines and the Guidelines of the European Union Council (63/2010/EU) following the Spanish regulation (RD53/2013) for the use of laboratory animals and were approved by the Scientific Committee of the University of Leon. All efforts were made to minimize animal suffering and to reduce the number of animals used.

### 2.2. Surgery and drug administration

Transient middle cerebral artery occlusion (tMCAO) was performed as previously described [21,34]. Briefly, anesthesia was induced with 3.5-4% isoflurane (Esteve, Barcelona, Spain) in 100% O_2_-enriched air with a flow of 2 L/min and maintained at 2% isoflurane during the surgery. The body temperature was maintained at 36 ± 1°C during surgery with a feedback-regulated heating pad monitored with a rectal probe. A Doppler probe (Perimed, Järfälla, Sweden) was fixed on the temporal bone over the MCA to monitor blood flow. Once the carotid bifurcation was exposed, a 4-0 silicon-coated monofilament (Doccol Corporation, Sharon, MA, USA) was inserted into the right common carotid artery and led through the right internal carotid artery until blocking of the origin of the MCA. Blood flow blockage was monitored by a Doppler probe, and animals with less than 80% of blood flow restriction were discarded. After one hour of occlusion, the monofilament was withdrawn, allowing blood reperfusion while monitoring by a Doppler probe and incisions were permanently sutured. The same surgery was performed in sham animals, except for MCA occlusion with the monofilament. No analgesic drugs were administered to avoid a bias in the interpretation of the data. The dose-response assay of meloxicam was performed administering of 0.5 mg/kg (Mel0.5), 1 mg/kg (Mel1), 5 mg/kg (Mel5), and 10 mg/kg (Mel10) of meloxicam (Boehringer Ingelheim, Ingelheim am Rhein, Germany) intravenously (i.v.) through the tail vein, one hour and 24 hours after reperfusion. Vehicle (saline) was administered to vehicle (Veh) and sham animals (Sham). For studies with a longer time of reperfusion (7 days), meloxicam 1 mg/kg was administered in the acute phase (1 h and 24 h after reperfusion, i.v.; Mel1-a) or as a chronic dose (one hour after reperfusion and then a repeated dose every 24 hours until sacrifice, i.v; Mel1-c).

### 2.3. Behavioral analysis

The neurological deficit was evaluated based on the modified Neurological Severity Score (mNSS) [35] at 24 h, 48 h, and 7 days after reperfusion. For this, rats were video recorded for 5 minutes on a behavioral table and then were evaluated by two blinded and independent observers. Results are shown as the average given by the two researchers at each time point.

The cylinder test was performed after 7 days of reperfusion. The rats were placed in a plastic cylinder (37.5 cm high 13 cm diameter) and filmed for 10 min. A blinded observer scored the first 20 forelimb contacts with the cylinder. The use of the contralateral over the ipsilateral forepaw was calculated as the bias on the contralateral using the following formula: bias (%) = 100 * (contralateral) / (ipsilateral + contralateral) [36].

### 2.4. Infarct volume

After 48 h of ischemia, rats were decapitated, and their brains were removed and placed in a cold rodent brain matrix (ASI Instruments, Warren, MI). Coronal 2 mm-thick sections were obtained from the bregma 3.2 to bregma -6.8. The coronal sections were incubated in 1% 2,3,5-triphenyl tetrazolium chloride (TTC) in 50 mM phosphate-buffered saline (PBS), pH 7.4 for 25 mins at 37°C. Then, sections were fixed in 4% paraformaldehyde (PFA) in PBS overnight. Brain sections were digitized at 600 dpi resolution (Canon Inc, Ohta-ku, Tokyo), and infarct volume was measured with ImageJ software (NIH). Infarct volume was calculated as: Infarct volume (%) = 100 x [non-stained volume (mm^3^)/ total volume (mm^3^)].

### 2.5. C reactive protein

Blood was collected at the moment of the and centrifuged immediately at 1200 xg for 10 min at 4°C. The serum was aliquoted and stored at -80°C until further use. C-reactive protein (CRP) was measured by an ELISA kit (cat. number CYT294, Millipore Sigma, Saint Louis, MO) following the manufacturer’s instructions.

### 2.6. Immunofluorescence and image acquisition

Rats were sacrificed with a sublethal dose of sodium pentobarbital (200 mg/kg) (Esteve, Barcelona, Spain) and were perfused via the aorta with 4% PFA. Brains were fixed, cryoprotected, and sectioned, as previously described [34]. Seven equidistant 40-micron coronal sections separated by 960 µm between bregma 2.20 mm and bregma -3.80 mm were used for neuronal nuclear protein (NeuN) staining. Three sections separated by 960 µm between bregma 1.24 and bregma -0.68 were used for ionized calcium-binding adapter molecule 1 (IBA-1), glial fibrillary acidic protein (GFAP), phosphorylated growth-associated protein 43 (pGAP43), and doublecortin (DCX) immunostainings. Primary and secondary antibodies are summarized in Table 1. A Zeiss LSM 800 confocal microscope was used to acquire images from the immunostained sections. Bias in the image quantification was avoided by keeping constant the pinhole, detector gain, laser power, and pixel dwell.

**Table 1.** Antibodies and specific conditions used in immunofluorescence assays. All tissues were exposed to an antigen retrieval solution (citrate buffer pH 6.0) for 25 min at 95°C, DAPI was used to contrast nuclei in immunofluorescence assays.

| Conditions for Immunofluorescence assays |  |  |  |
| --- | --- | --- | --- |
| Primary antibody | Blocking agent | Incubation solution | Secondary antibody |
| Goat anti IBA-1, 0.25 µg/mL<br>(ref: Ab5076, Abcam, Cambridge, MA, USA) | BSA 1% | Triton 0.2%<br>in PBS | Alexa fluor 647 donkey anti goat, 1 µg/mL<br>(ref: A21448, Thermo Fisher Scientific, Waltham, MA USA) |
| Rabbit anti GFAP, 6 µg/mL<br>(ref: ZO334, Dako, Santa Clara, CA, USA) | Goat serum 20% | Triton X100 0.2%<br>in PBS | Alexa fluor 488 goat anti rabbit, 1 µg/mL<br>(ref: A11055, Thermo Fisher Scientific, Waltham, MA USA) |
| Mouse anti NeuN, 2 µg/mL<br>(ref: MAB337, Millipore, Burlington, MA, USA) | Goat serum 20% | Triton X100 0.2%<br>in PBS | Alexa fluor 568 goat anti mouse, 1 µg/mL<br>(ref: A21069, Thermo Fisher Scientific, Waltham, MA USA) |
| Rabbit anti DCX, 2 µg/mL<br>(ref: 4604, Cell Signaling Technology Danvers, MA, USA) | Goat serum 20% | Triton X100 0.2%<br>in PBS | Alexa fluor 488 goat anti rabbit, 1 µg/mL<br>(ref: A11055, Thermo Fisher Scientific, Waltham, MA USA) |
| Rabbit anti pGAP43, 8 µg/mL<br>(ref: Ab194929, Abcam, Cambridge, MA, USA) | Goat serum 20% | Triton X100 0.5%<br>in PBS | Alexa fluor 488 goat anti rabbit, 1 µg/mL<br>(ref: A11055, Thermo Fisher Scientific, Waltham, MA USA) |

#### 2.6.1. Microglia activation analysis

A modification of the optical dissector method [37] was used to count IBA1+ cells and perform morphometric fluorescent and densitometric analyses. In brief, a grid of 0.255 mm^2^ squares with a lateral resolution of 0.156 µm/pixel was acquired with a Plan-Apochromat 40×/1.3Oil DIC (UV) VIS-IR M27. In each dissector, Z-stack images were taken separated by 4 µm along the Z-axis (5 images, 20 µm in total) (see Figure 1H). IBA-1+ cells were counted in each dissector, and the average of the dissectors in each rat was expressed as the number of IBA-1+ cells/mm^3^. The degree of microglial activation was estimated by measuring the number of branches and the length of the cell processes as previously described [38]. Briefly, each dissector was binarized, skeletonized, and analyzed with Analyze Skeleton 2D/3D plugin in ImageJ, and the results were normalized with the number of IBA-1+ cells in each dissector and expressed as endpoints/cell and the summed process length/cell.

**Figure 1.**
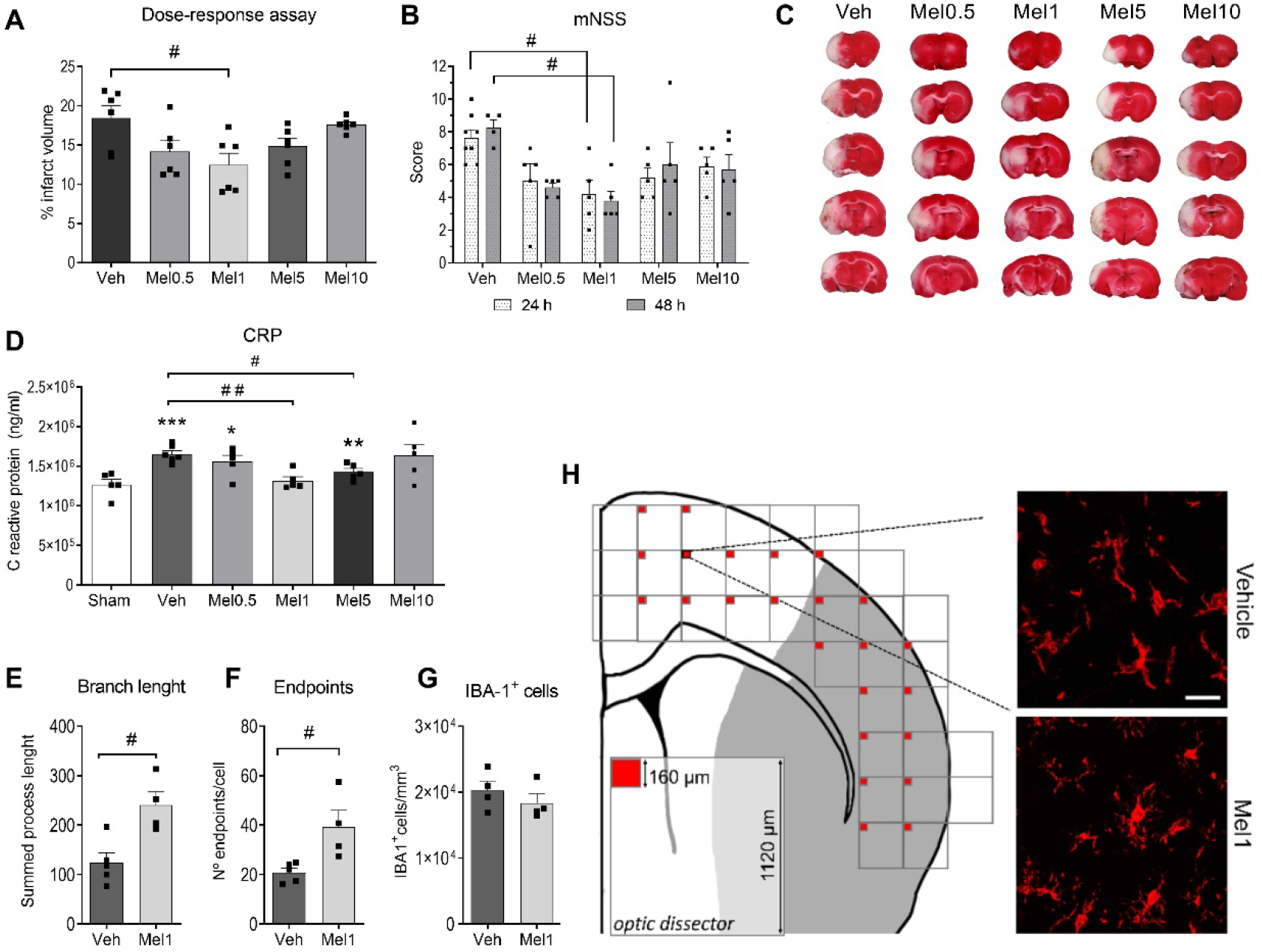
Neuroprotective and anti-inflammatory effects of meloxicam after 48 h of reperfusion. A) Percentage of infarct volume in the dose-response assay after 48 h of reperfusion. B) Neurological score measured at both 24 and 48 hours after reperfusion. C) Representative brain slices stained with TTC. D) CRP levels in plasma and microglial activation measured as E) branch length, F) number of endpoints, and G) number of IBA-1+ cells. H) Schematic representation of image acquisition (red squares) by optical dissector method to measure microglial activation in the cerebral cortex and representative images of IBA-1+ cells. One-way ANOVA followed by the Tukey test in the TTC assay and CRP assay and followed by the Kruskal-Wallis test for the behavioral assay. Student’s t-test was used for the microglial activity assay. # represents significant differences with vehicle animals and * with sham animals. ANOVA: Analysis of variances; CRP: C reactive protein; IBA-1: ionized calcium-binding adapter molecule; mNSS: modified neurological severity score; TTC: 2,3,5-triphenyl tetrazolium chloride.

#### 2.6.2. MNeuronal loss quantification

For NeuN staining, each section was scanned to obtain a high-resolution image (1.25 µm/pixel) of the whole section using the Tile Scan Module (included in Zen Blue software) with a Plan-Apochromat 10×/0.45 M27 objective. Some parts of the ischemic area came off due to the strong damage in the tissue, thus to avoid bias, the fluorescence image was overlapped with the corresponding coronal section of the Patxinos rat brain atlas (reference). The limit of the area considered as “neuronal loss” is the lack of clear and defined neuronal bodies, as shown in the insets in Figure 2E. Then, neuronal loss was quantified on the tile sections with the Image J software. Results were expressed as Percentage of neuronal loss = 100 × (non-stained area / total analyzed area).

**Figure 2.**
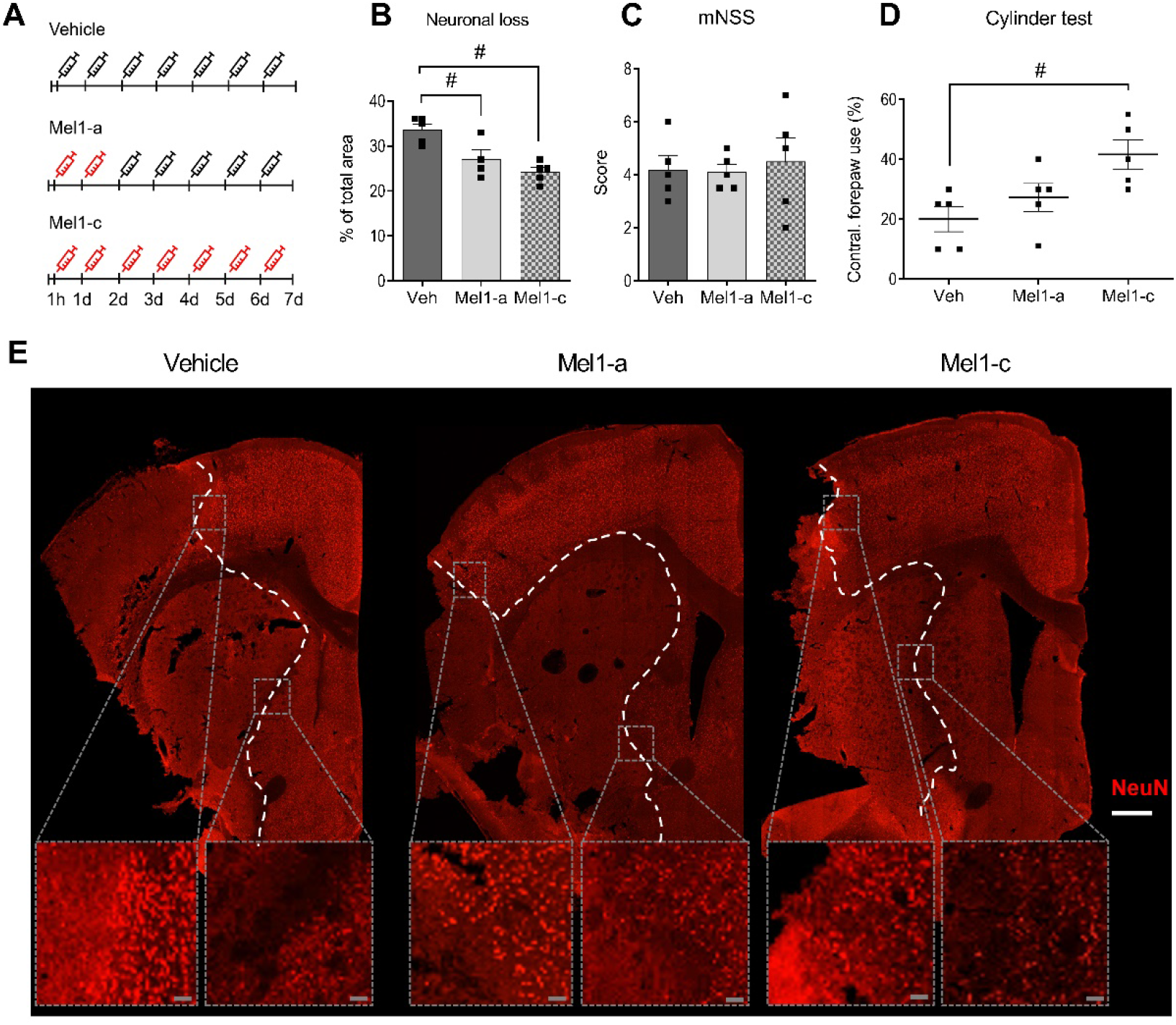
Long-term neuroprotection of meloxicam treatment. **A) Schematic representation of meloxicam dosage at 7 days of reperfusion**. Red syringes represent a meloxicam administration of 1 mg/kg and white syringes represent vehicle administration. B) Percentage of neuronal loss after 7 days of reperfusion measured as the area without NeuN staining. Behavioral analysis was assayed with C) mNSS and D) the cylinder test. E) Representative images of NeuN staining. Inset shows in detail the limit considered in this study to measure the neuronal loss. White scale bar: 200 μm; Gray scale bar: 20 μm. # represents significance with vehicle condition. One-way ANOVA followed by Tukey’s test was performed to analyze neuronal loss and Kruskal–Wallis to analyze the behavioral tests. mNSS: modified Neurological Severity Score; NeuN: neuronal nuclear protein.

#### 2.6.3. Glial scar analysis

Z-stack images were acquired separated by 320 µm between them (distance between the center of each image), setting the first image in the edge of the glial scar (see Figure 3A). During the analysis, GFAP labeling was used to create a binary mask applied to GFAP images to delimitate the shape of the astrocytes. The total fluorescence intensity (TFI) was obtained in order to normalize the total relative amount of GFAP per cell. TFI values were normalized using the number of astrocytes in each dissector and the average TFI for each rat was expressed as TFI/cell.

**Figure 3.**
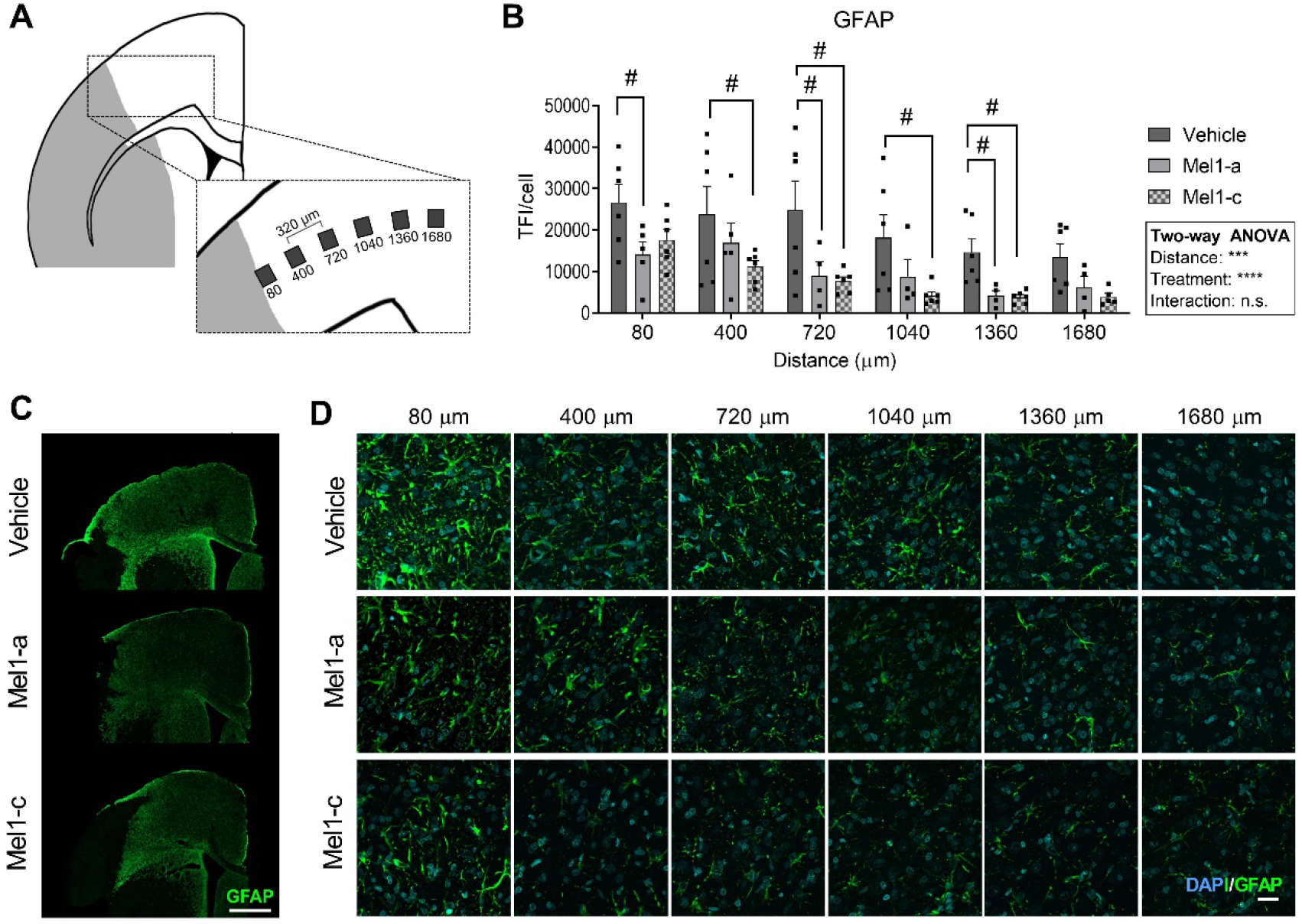
Astrocyte reactivity in the glial scar in the cerebral cortex. **A) Representative scheme of the specific analyzed regions in the glial scar**. B) GFAP TFI/cell at each point of analysis along the glial scar. Representative images of GFAP immunostaining are shown as C) a whole image of the cerebral cortex and D) in each analyzed region. Scale bar: C) 200 μm and D) 20 μm. Two-way ANOVA was performed, followed by Tukey’s test. # represents significance with vehicle condition. GFAP: glial fibrillary acidic protein; TFI: total fluorescence intensity.

#### 2.6.4. Axonal sprouting analysis

Z-stack images were obtained on the edge of the ischemic damage. This immunostaining presents sparse labeling due to the relatively low amount of this protein and high background due to the proximity to the damage. For this reason, we first produced a single image with the sum of fluorescence of all the z-stack. Then, immunolabeling with pGAP43 was used to create a mask based on an intensity threshold to measure the immunolabeling signal accurately without considering the background. The parameters of the threshold were maintained constant in all the analyzed images to avoid bias. Mean fluorescence intensity (MFI) was measured as a representation of the average relative protein amount in each analyzed region. The average of the MFI for each rat in the region of interest (shown in Figure 4A) was expressed as MFI/mm^2^.

**Figure 4.**
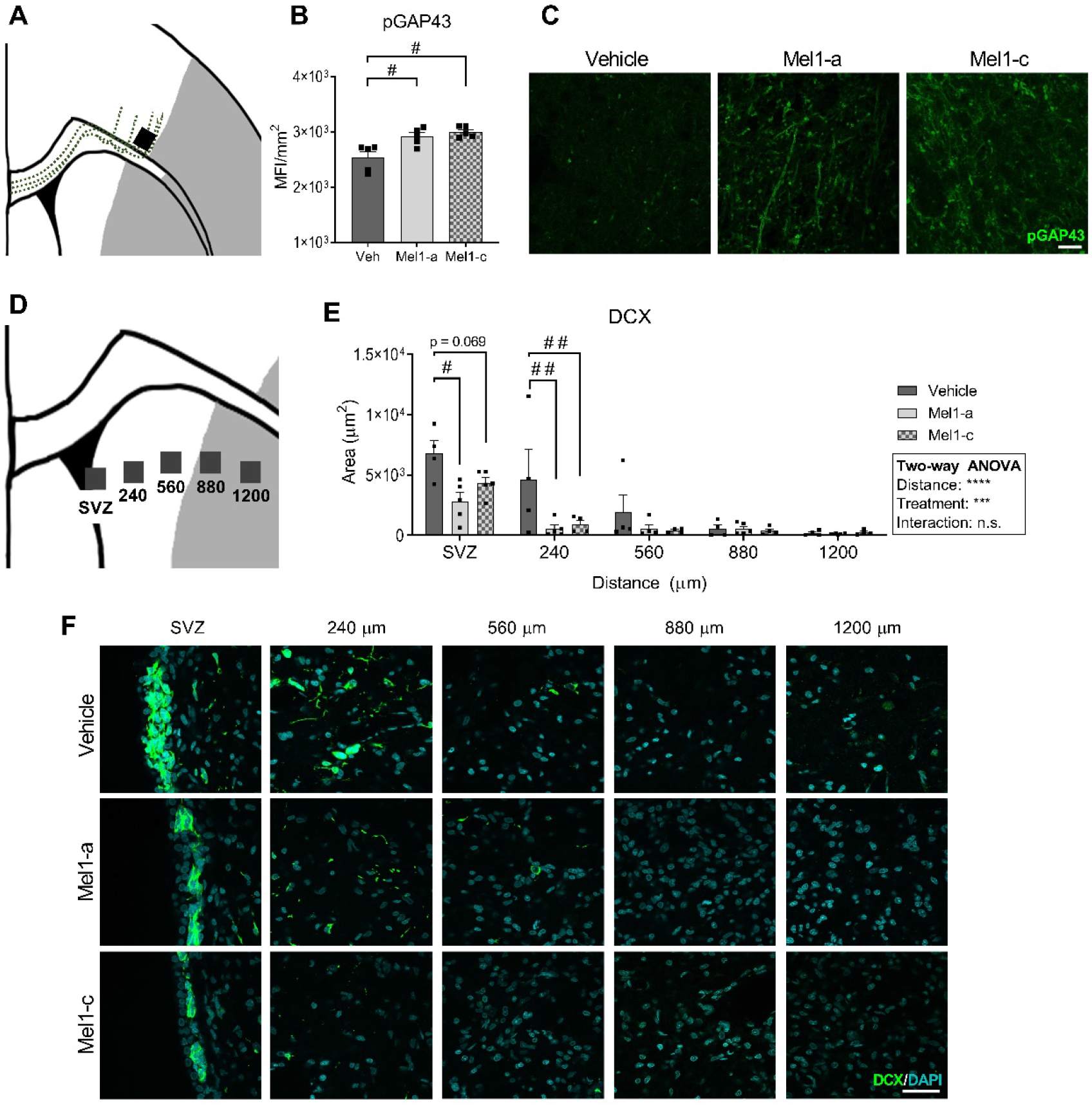
Regenerative events after 7 days of reperfusion. A) Scheme showing the analyzed regions (gray squares) in the pGAP43 immunostaining. Dotted line represents one of the possible pathways described for axonal sprouting. B) Protein levels of pGAP43 expressed as the mean fluorescence intensity per mm^2^ and C) representative images of the pGAP43 immunostaining (scale bar: 20 μm). D) Scheme showing the analyzed regions (gray squares) in the DCX immunostaining. E) Area occupied by DCX+ cells in the different analyzed regions and F) the representative images of each analyzed region (scale bar: 20 μm). One-way ANOVA followed by Tukey’s test was performed to analyze pGAP43 and two-way ANOVA followed by Tukey’s test was performed to analyze DCX. # represents differences with respect to vehicle condition. DCX: doublecortin; MFI: mean fluorescence intensity; pGAP43: phosphorylated growth-associated protein 43; SVZ: subventricular zone.

#### 2.6.5. Neuronal progenitor analysis

Z-stack images were obtained from the subventricular zone (SVZ), as well as different regions of the striatum located perpendicular to the SVZ were imaged (shown in Figure 4D). Due to the impossibility of discerning individual cells in the SVZ, the area stained with DCX was measured with ImageJ in each dissector. The average values per rat in each analyzed region are expressed as DCX+ area/mm^3^.

### 2.7. Statistical analysis

Statistical analyses were carried out using GraphPad Prism 6 (GraphPad software). One-way ANOVA followed by Tukey’s test was performed to analyze the differences between treatments in parametric datasets. Two-ways ANOVA followed by Tukey’s post hoc was performed to analyze the effect of the treatments in different areas. As values obtained from the behavioral test are considered non-parametric, the Kruskal-Wallis followed by the Dunn’s test was performed to analyze them. Data are presented as mean ± SEM. The number of samples (n) differs between assays due to additional difficulties in analyzing several parameters in extremely damaged tissue, thus each dot represents one animal. Significance was set at p < 0.05. One, two, three and four significant symbols refer to p values of p < 0.05, p < 0.01, p < 0.001 and p < 0.0001, respectively.

## 3. Results

### 3.1. Meloxicam reduces the infarct volume and improves post-ischemic behavior after 48 h of reperfusion

To set up the optimal neuroprotective dose of meloxicam, a dose-response assay measuring the infarct volume after 48 h of reperfusion was performed. The different dosage displayed a U-shaped curve where the neuroprotective effects reached statistical significance at 1 mg/kg of meloxicam (Mel1). Consistently, behavioral data at both 24 h and 48 h after reperfusion only displayed significant improvement for 1 mg/kg of meloxicam (Figure 1B).

After 48 h of reperfusion, the levels of CRP were used to measure systemic inflammation in plasma. The range of doses assayed showed a U-shaped curve and only animals treated with 1 mg/kg meloxicam reached significantly lower CRP levels than vehicle animals (Figure 1D). To corroborate the anti-inflammatory effect, the microglia activation in the ischemic hemisphere (ipsilateral) was only measured after the administration of the neuroprotective dose (1 mg/kg of meloxicam). The total branch length and endpoints of IBA-1+ cells (Figures 1E and 1F, respectively) significantly increased in treated animals. No changes in the number of these cells were detected (Figure 1G).

### 3.2. Neuroprotection remains after 7 days of reperfusion

To determine if neuroprotection exerted by 1 mg/kg is maintained after 7 days of reperfusion, we measured the neuronal demise after acute (Mel1-a) and chronic (Mel1-c) treatments with meloxicam (Figure 2A,E). Neuronal loss observed in vehicle animals was decreased by the acute and chronic treatments with meloxicam (Figure 2B). Acute and chronic treatments presented similar results. The behavior analysis using the mNSS did not reveal differences between vehicle and treated animals (Figure 2C). However, animals chronically treated with meloxicam displayed improved motor behavior in the cylinder test compared to vehicle animals (Figure 2D).

### 3.3. The reactivity of the astrocytes in the glial scar is modulated by both chronic and acute doses of meloxicam

Astrocytes at the edge of cortical glial scar showed a hypertrophic morphology characterized by engrossed cell bodies with wide branches, which were observed at 80 and 400 μm from the border and were progressively replaced by less hypertrophied cells further from the edge (Figure 3D). The reactivity of astrocytes in the glial scar was quantified as the levels of GFAP per cell (TFI/cell). This analysis revealed a distance-dependent decrease in the astrocyte reactivity, attenuated by both acute and chronic meloxicam doses in each analyzed point (Figure 3B).

### 3.4. Meloxicam treatment enhances axonal sprouting after 7 days of reperfusion

Axonal sprouting was measured in the edge of the glial scar as a molecular parameter of the tissue’s new connectivity attempt (Figure 4A) using pGAP43, a specific marker for axonal sprouting [39]. The analysis revealed that both chronic and acute treatment with meloxicam significantly increased the levels of pGAP43 after 7 days of reperfusion (Figure 4B,C).

### 3.5. Stroke-induced neurogenesis after 7 days is decreased by meloxicam

To analyze the effect of meloxicam in neurogenesis, we labeled the neuronal progenitor cells (NPCs) with DCX and analyzed them in both the SVZ (NPCs) and in the striatum (migrating NPCs) (Figure 4D). The area occupied by DCX cells in the SVZ was significantly lower in animals treated with an acute dose of meloxicam compared to vehicle animals. Chronic treatment with meloxicam also decreased the area of DCX cells in the SVZ, but we failed in finding significance. The closest region to the SVZ analyzed also displayed significant reductions in meloxicam-treated animals (Figure 4E). No differences were observed in the different conditions at 560, 880 and 1200 µm of distance from the SVZ (Figure 4C).

## 4. Discussion

### 4.1. The neuroprotective effect of meloxicam: more is not always better

Previous studies suggest that meloxicam could be used as a potential treatment for stroke [31,33,40]. This led us to analyze its potential palliative effect in a model of transient focal ischemia. The optimal dose set-up is an indispensable step in this study since the balance of inflammation after stroke may worsen the damage or provide a null effect in terms of neuroprotection. Our dose-response analysis revealed a U-shaped response in a short range of doses (from 0.5 mg/kg to 10 mg/kg). Our results fit previous reports addressing a loss of COX-2 selectivity when meloxicam is administered at high doses [22,23,41]. Thus, meloxicam seems to represent one more example of hormesis, a concept that includes the inverted U- or U-shaped curves in dose-response studies. This phenomenon has been explained as a limited, temporally based overcompensation after a disruption in homeostasis and has been predicted to be near to universality, although it is not observed in the majority of cases, and represents an evolutionary strategy to select biological optimization responses [42]. Since meloxicam is a preferential COX-2 inhibitor, its hormetic effect should be mediated through the COX-2 pathway. However, it is likely that the curve-response of meloxicam is also modulated by a disbalance in the selective inhibition by meloxicam through a higher COX-1:COX-2 ratio, leading to gastrointestinal damage and hepatotoxicity, effects that have been linked to an overdose of meloxicam [43,44]. The lack of systemic anti-inflammatory effect in the higher doses assayed in this stroke model makes it difficult to choose an adequate dosage to obtain significant effects, which perhaps explains the low number of references on the use of meloxicam in stroke models. However, when the proper dosage is characterized, the treatment results in a very consistent response after 48 h of reperfusion both in the reduction of infarct volume and improvement of the neurological deficit. In fact, meloxicam presents similar outcomes to those observed with a selective anti-COX-2 agent, celecoxib, the only *coxib* with clear neuroprotective effects due to its unique properties not shared with other *coxib* agents [34,45].

### 4.2. Acute treatment with meloxicam is enough to maintain neuroprotection and modulate the glial scar reactivity after 7 days of reperfusion

Neuroprotective effects similar to those observed after 48 hours were maintained after 7 days when animals received either acute or chronic treatment with meloxicam. These data show that the two first doses of meloxicam are enough to exert a neuroprotective effect maintained over time. However, only animals treated with the chronic dose presented significantly improved behavior, while animals treated with the acute dose only present a tendency to increase. This improvement after the chronic administration could partially rely on meloxicam’s ability to reduce post-surgery pain, thus improving the animals’ performance in the cylinder test. However, there was a clear tendency of improvement related to the neuroprotective effect of meloxicam with the acute dose, even when it was not statistically significant. The mNSS is only suitable for short reperfusion times because the parameters of evaluation in this test have almost disappeared at that time point.

Astrocyte involvement in the local inflammatory response after ischemia is widely described [46–48]. The formation of the glial scar limits the extent of the damage, but also interferes with the regenerative response attempts by secreting inhibitory factors [49,50]. The lack of differences in the glial scar reactivity between the acute and chronic administrations of meloxicam suggests that the main signals involved in glial scar formation and astrocyte reactivity are triggered in the first 48 hours. These data match with the time course of two signaling pathways involved in astrocyte hypertrophy, i.e. GFAP overexpression and glial scar formation: the nuclear factor kappa B (NF-κB) pathway and signal transducer and activator of transcription (STAT3) [11,51]. NF-κB signaling is activated in the first hours after reperfusion [52], mainly by pro-inflammatory cytokines [53], leading to the release of interleukin 6 (IL-6), which activates STAT3 [54]. STAT3 activation is essential to GFAP overexpression and glial scar formation [55]. STAT3 is activated 2-4 h after reperfusion [56], and it remains activated for 24 h later [57]. These data suggest that inflammation in the first hours after a stroke is crucial to the initiation of astrocyte reactivity and glial scar formation and would explain why acute and chronic meloxicam are similarly effective in modulating astrocyte reactivity in the cerebral cortex. However, more deep studies focused on the molecular aspects of inflammation should be carried out to elucidate this specific mechanism.

### 4.3. The modulation of inflammation affects the regenerative processes after stroke in divergent ways

Regeneration after stroke is triggered in many different ways, including new cell generation and the induction of synaptic plasticity [58]. These two mechanisms can be independently activated depending on the model, the reperfusion timing, and the extrinsic conditions or treatments. Our results show that the modulation of inflammation with meloxicam increases axonal sprouting, but reduces new neuron generation in the SVZ. The basis underlying the increase in axonal sprouting could be related to the reduction of astrocyte reactivity. Axonal sprouting is part of the tissue reorganization and scar formation [59]. Complete ablation of glial scar prevents axonal outgrowth, suggesting that the presence of a glial scar and astrocyte reactivity are necessary to axonal regeneration [49,50]; however, excessive astrocyte activation could lead to the production of inhibitors of axonal outgrowth [51,60]. In this study, the modulation of the glial scar by meloxicam seemed to promote axonal sprouting at the edge of the glial scar, which can be easily attributed to an attempt of the tissue to promote the regeneration of lost connections.

Besides its positive effect in axonal sprouting, meloxicam decreases the generation of new neurons in the SVZ. The considerable variability in the data obtained seems to be responsible for the lack of significant differences between the meloxicam chronic treatment dose compared to vehicle treated animals, probably due to the inherent variability in the extent of the infarct volume of this model. However, the observed tendency suggests that this mechanism is also elicited in the first 48 hours since both acute and chronic treatments decrease NPC formation. This effect has also been reported in non-pathological conditions in both the SVZ and the subgranular zone (SGZ) in healthy animals treated with meloxicam [61], and similar effects have been reported in COX-2 KO mice [62]. The influence of COX inhibition on neurogenesis is not fully understood, but it is related to the decrease of prostaglandins such as prostaglandin E2 (PGE2). PGE2 transactivates a receptor for the ependymal growth factor (EGF) triggering the mitotic signaling [63]. Thus, inhibition of PGE2 by anti-inflammatory agents could account for the decreased mitotic signaling and a reduction in the division of NPCs [62]. The strong effect of meloxicam on inhibiting PGE2 production [64] could explain these results. Another possible mechanism could rely on microglia since microglia-derived factors stimulate the early stages of neurogenesis and promote NPC recruitment to the sites of inflammation [65,66]. In this sense, the reduction of microglial reactivity by meloxicam observed after 48 h of reperfusion could prevent NPC promotion. Other mechanisms could rely on gamma-aminobutyric acid (GABA). GABAergic signaling blocks cell proliferation, while GABAergic inhibition increases NPC proliferation and differentiation [67]. A study carried out in a hippocampal slice culture model of ischemia showed that meloxicam promotes the expression of GABA_A_ receptors [31], suggesting a putative role of meloxicam in increasing GABAergic signaling. All these mechanisms are plausible and may participate in the inhibition of neurogenesis after meloxicam treatment. Whether this inhibition is detrimental or irrelevant to the recovery after stroke is a matter that should be analyzed in further studies accounting for longer times of reperfusion and more specific behavior analysis.

The surprising similarities in the response after acute and chronic administration of meloxicam support the idea that the short-term treatment of inflammation is of special relevance. Moreover, it shows that the modulation of inflammation in early timepoints has repercussions on later events of regeneration processes, such as axonal sprouting and neurogenesis. This modulation presents special relevance in the search for putative therapies for stroke since many studies are focused only on providing neuroprotection by reducing the infarct volume and they do not consider the effects on regenerative attempts. The tight connection between inflammation and regeneration makes it imperative to study these events after experimental anti-inflammatory treatments.

## 5. Conclusions

This study shows for the first time that meloxicam exerts neuroprotection in a tMCAO model and how this neuroprotective effect is linked to both the promotion and the reduction of different regenerative events that occur after stroke. In summary, this study provides a perspective analysis of the use of anti-inflammatory treatments after stroke and how these treatments should be carefully analyzed in the short and long term after stroke.

## Author contribution

Irene F Ugidos: Conceptualization, Investigation, Formal Analysis, Writing-Original Draft; Paloma González-Rodríguez: Investigation; Maria Santos-Galdiano: Methodology, Investigation; Enrique Font-Belmonte: Investigation; Berta Anuncibay-Soto: Project Administration; Diego Pérez-Rodríguez: Conceptualization, Software, Supervision; Arsenio Fernández-López: Supervision, Funding acquisition, Writing-Review and Editing.

## Acknowledgements

This work was supported by MINECO and FEDER funds (RTC-2015-4094-1), by Junta de Castilla y León (LE025P17) and by Neural Therapies SL (NT-Dev-01).

